# DIVERSITY OF GENOME SIZE AND CHROMOSOME NUMBER IN HOMOTHALLIC AND HETEROTHALLIC STRAINS OF THE *CLOSTERIUM PERACEROSUM–STRIGOSUM–LITTORALE* COMPLEX (DESMIDIALES, ZYGNEMATOPHYCEAE, STREPTOPHYTA)

**DOI:** 10.1101/2023.05.01.538656

**Authors:** Yuki Tsuchikane, Misaki Watanabe, Yawako W Kawaguchi, Koichi Uehara, Tomoaki Nishiyama, Hiroyuki Sekimoto, Takashi Tsuchimatsu

## Abstract

Members of the *Closterium peracerosum–strigosum–littorale* (*C. psl.*) complex are unicellular zygnematophycean algae, which are suggested to be closely related to land plants. A zygospore is typically formed as a result of conjugation between mating-type plus (mt^+^) and mating-type minus (mt^−^) cells during sexual reproduction in heterothallic strains. On the other hand, zygospores are formed between genetically identical cells in homothallic strains. In this study, we isolated novel homothallic strains in the *C. psl.* complex. Phylogenetic analysis revealed the polyphyly of homothallic strains, suggesting multiple transitions between homothallism and heterothallism in the *C. psl.* complex. We measured the 1C genome size of the *C. psl.* complex by using flow cytometry after staining nuclei with propidium iodide, which ranged from 0.53 to 1.42 Gbp. We counted chromosome numbers using confocal microscope images, finding that two homothallic strains had fewer chromosomes than four heterothallic strains. Genome size positively correlated with both the cell size and chromosome number. Chromosome numbers differed even within the same mating group, suggesting a mechanism tolerating chromosomal rearrangements during meiosis in the *C. psl.* complex.

## INTRODUCTION

The evolutionary transition of mating systems between outcrossing (self-sterility) and self-fertilization (selfing) is frequently observed in plants, fungi, as well as in animals (Wright et al. 2013, Shimizu and Tsuchimatsu 2015, Cutter 2019). The transitions of mating systems are often accompanied by various changes in reproductive, life-history, genomic, and cytological traits (Wright et al. 2013, Shimizu and Tsuchimatsu 2015, Cutter 2019, Tsuchimatsu and Fujii 2022). For example, selfing species tend to show lower nucleotide diversity and effective population size compared with outcrossing relatives, and accumulate more slightly deleterious mutations, including chromosomal rearrangements (Coyne and Orr 2004, Wright et al. 2013). Mating system transitions are also observed in haploid organisms that are mostly isogamous (e.g., Hanschen et al. 2018, Yamamoto et al. 2021). While gametes from a single clone conjugate with each other and produce viable progenies in homothallic (self-fertile) organisms, gametes from two different genetic clones are required for successful mating in heterothallic (self-sterile) organisms (Graham and Wilcox 2000). Such changes in association with mating system transitions are well documented in multicellular organisms, but relatively unexplored in unicellular organisms, despite their diversity in mating systems.

Members of the *Closterium peracerosum–strigosum–littorale* (*C. psl.*) complex are unicellular isogamous algae in Zygnematophyceae, which are the sister clade of land plants (Jiao et al. 2020, Wickett et al. 2014). In the *C. psl.* complex, both heterothallic and homothallic strains have been reported (Tsuchikane and Sekimoto 2019). Heterothallic strains consist of two sexes: mating-type plus (mt^+^) and mating-type minus (mt^−^). The details of sexual reproduction have been studied from physiological, biochemical, and molecular biology perspectives (Abe et al. 2011, Kanda et al. 2017, Kawai et al. 2022, Sekimoto 2017, Sekimoto et al. 2012). Recently, genomes of the two mating types (NIES-67 and NIES-68) were sequenced (Sekimoto et al. 2023). When mt^+^ and mt^−^ cells were mixed in a nitrogen-depleted medium in the presence of light, they divided once to form sexually competent gametangial cells through a process called sexual cell division (Ichimura 1971). Two gametangial cells of different mating types form paired cells, which release their gametic protoplasts and fuse immediately to form a zygospore. The conjugation process of the homothallic strain has also been reported (Tsuchikane et al. 2010). The first step in the conjugation process is mitotic cell division, which results in the formation of two sister gametangial cells from a vegetative cell. These two gametangial cells form a sexual pair followed by a zygospore. Approximately 90% of homothallic zygospores originate from the conjugation of two sister gametangial cells, derived from one vegetative cell (Tsuchikane et al. 2010).

The pattern of reproductive isolation and mating systems have been studied together with the phylogenetic relationship in the *C. psl* complex. At least five reproductively isolated heterothallic mating groups have been reported (groups II-A, II-B, II-C, I-E, and G) (Kobayashi et al. 2022, Tsuchikane et al. 2018b, Watanabe 1977, Watanabe and Ichimura 1978). Although zygospore formation is restricted between strains of different mating groups, a few zygospores are formed between mating groups II-A and II-B, indicating incomplete reproductive isolation (Watanabe and Ichimura 1978, Tsuchikane et al. 2008). To date, only a single clonal culture for the homothallic strain (kodama20: NIES-2666) has been established. This homothallic strain was suggested to form a phylogenetically monophyletic unit with the heterothallic mating group II-B (Tsuchikane et al. 2012), and hybrid zygotes can be formed between them (Tsuchikane et al. 2012). More strains, in particular homothallic strains, would be required to understand the process of mating system transitions with accompanied cytological and genomic changes in the *C. psl* complex.

Several studies have been conducted on cytology in zygnematophycean algae (Godward 1966, Poulíèková et al. 2014, Mazalová et al. 2011). In *Closterium*, the number of chromosomes was estimated in *Closterium ehrenbergii* Meneghini: 100–106 for mating group A, 97–104 for mating group B, and 105–127 for mating group C (Kasai and Ichimura 1984). Flow cytometry has often been used to estimate genome size in flowering plants (Doležel et al. 1992), but, to our knowledge, it has not been performed in any species of *Closterium*.

In this study, we first isolated homothallic and heterothallic strains of the *C. psl.* complex. Phylogenetic analysis revealed the polyphyly of homothallic strains, suggesting multiple transitions between homothallism and heterothallism in the *C. psl.* complex. We found extensive variation both in genome size (0.53–1.42 Gbp) and chromosome numbers (26.6±1.31 to 87.7±5.55) between strains. Genome size and chromosome number were correlated positively, but some of the homothallic strains showed large chromosome numbers with respect to genome size. This result suggests that each chromosome is smaller and more fragmented in these strains, possibly reflecting reduced constraints for structural rearrangements in homothallic strains. The *C. psl*. complex would serve as a new model to study cytological and genomic changes associated with mating systems.

## MATERIALS AND METHODS

### Strains and clonal culture conditions

The *Closterium* strains used in the present study are listed in Table 1. We obtained the following strains of the *C. psl*. complex from the National Institute for Environmental Studies, Environmental Agency (Ibaraki, Japan): NIES-53, 54, 58, 59, 64, 65, 67, 68, 261, 2666, 4319, 4320, 4321, 4322, 4581, and 4582. Strains Izu12-4-1, Hana, Naga37s-1, Shima19-9, Oki23-2, Biwa5-3, Izu12-7-4, Mie54-1, Miya184-1, Naga56-6, Oki21-8, Yama58-4, and Ho36-3 were established as follows. Water samples were collected from each locality using a plankton net (mesh size: 32 μm, RIGO, Tokyo, Japan) or disposable dropper (No. 4 dropper, Eiken Chemical Co. Ltd., Tokyo, Japan) using a previously reported method (Tsuchikane et al. 2018a). The soil sample from Naga37s-1 was collected from a paddy field in Nagano, Japan and air-dried. The dried soil sample (0.3–0.5 g) was placed in a petri dish (Φ 90 × 15). Sterile distilled water was poured into a petri dish to a height of 5 mm from the bottom. The petri dish was kept in an incubator at 23°C under a 16 h light/8 h dark cycle for approximately one week to germinate algal zygospores from the soil samples (dry soil method, Starr 1973). Light from LED lamps (PF20-S9WT8-D1; Nippon Medical and Chemical Instruments Co. Ltd. Osaka, Japan) was adjusted to 128 μmol photons·m^−2^·s^−1^ on the surface of the culture dish.

**Table 1.**
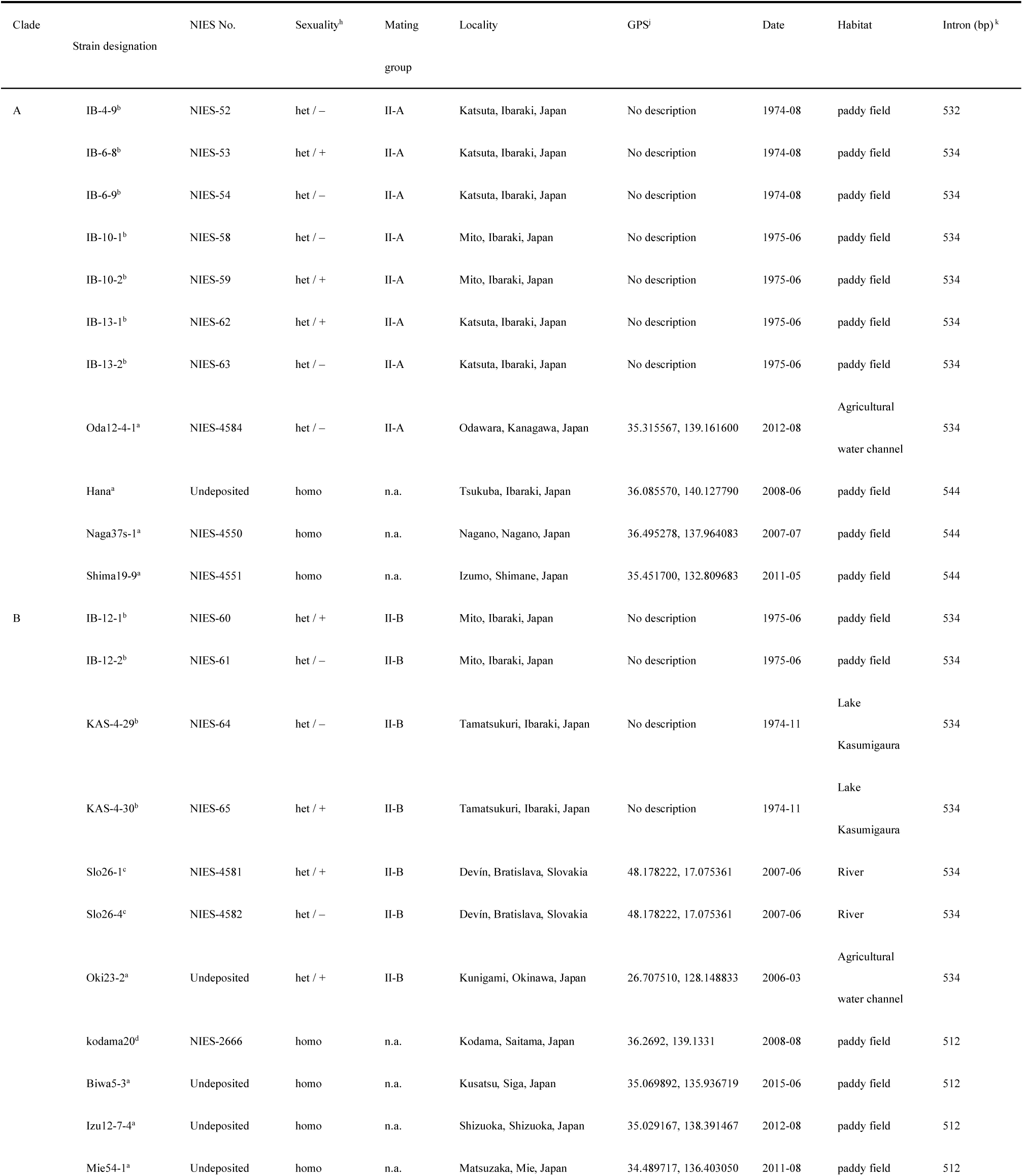
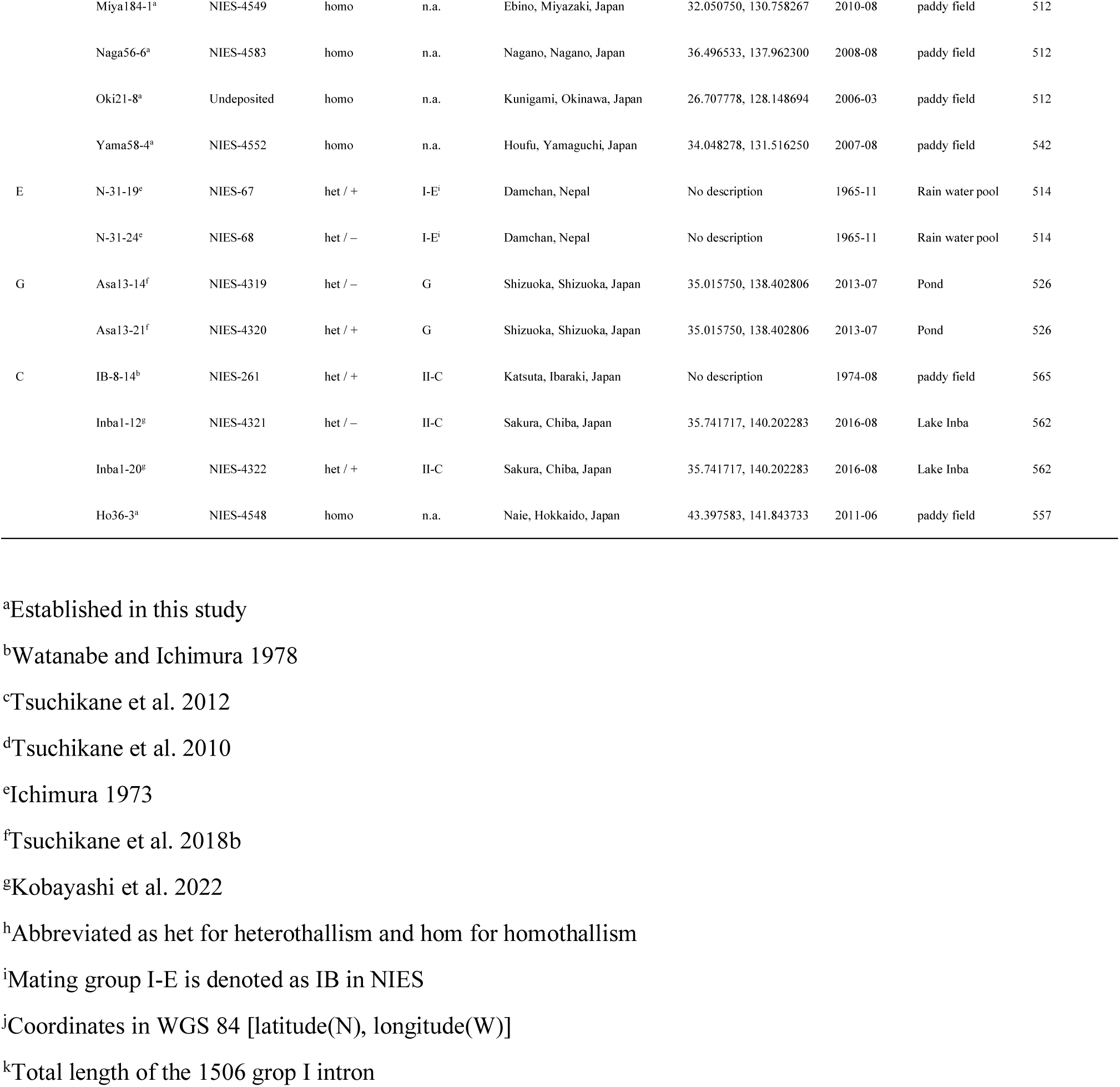
List of *Closterium* strains used in this study.

To proliferate isolated single cells, conditioned medium was prepared as follows. NIES-67 cells were cultured in nitrogen-supplemented medium (CA medium; Kasai et al. 2004) for 14–20 days. Subsequently, the medium was collected by filtration using filter paper (Advantec Toyo, Tokyo, Japan) and autoclaved (121°C for 20 min). The unknown factors for cell proliferation, which are secreted from growing cells into the surrounding environment, were included in the conditioned medium to facilitate cell division (Abe *et al*., 2011).

Cells morphologically identified as the *C. psl*. complex from water and the incubated soil samples were subsequently washed, isolated from the sample using the pipette-washing method (Pringsheim 1946, Tsuchikane et al. 2018a), and cultured in a test tube (18 mm diameter and 150 mm length) containing 16 mL of conditioned CA medium under the same culture conditions described above.

Proliferating clonal cells were inoculated into a 300 mL Erlenmeyer flask (AGC Techno glass co., Ltd. Shizuoka, Japan) containing 150 mL of CA medium under the same culture conditions described above. Vegetative cells in the late logarithmic phase (at 14 days) were collected by centrifugation (1,450×*g*, 5 min) and washed thrice with nitrogen-depleted mating medium (MI medium; Ichimura 1971). Ten milliliters of the cell suspension were prepared, and 100 μL of the cell suspension was diluted 100-fold. The cells contained in the diluted cell suspension were counted using a microscope equipped with a cell counter (Optical Plastic Plankton Counter, Matsunami Glass Ind., Ltd. Osaka, Japan) to determine the cell density of the undiluted cell suspension. A cell suspension that was assumed to contain 10,000 cells each was added, and the total cell number was adjusted to approximately 2 × 10^4^ in 2 mL of MI medium in a 24-well microplate (16 mm diameter wells; Iwaki Micro plate, Chiba, Japan).

Each heterothallic subclone was test-crossed with NIES-53 (mt^+^ of mating group II-A), NIES-54 (mt^−^ of mating group II-A), NIES-65 (mt^+^ of mating group II-B), and NIES-64 (mt^−^ of mating group II-B) to determine the mating type and mating group in a 24-well microplate (Iwaki). The cells were mixed and incubated under continuous light for 72 h. After incubation, zygospores were counted using a hemocytometer. The relative number of zygospores was calculated using the following equation:

Relative number of zygospores (%) = (number of zygospores × 2 / total number of cells) × 100.

Experiments were performed at least twice to confirm the reproducibility.

### Phylogenetic analysis

Vegetative cells in the late-logarithmic phase (at 14 days) were collected by centrifugation (1,450×*g*, 5 min) and counted. Thirty thousand cells of each clone were added to 100 μL of Quick Extract™ DNA Extraction Solution (Lucigen, Wisconsin, USA). DNA was extracted by heat treatment (65°C for 6 min and 98°C for 2 min). The 1506 group I intron interrupting the nuclear small subunit rRNA genes was amplified and directly sequenced using the protocols described by Tsuchikane et al. (2010). For phylogenetic analyses, identical sequences were treated as a single operational taxonomic unit. The sequences of the *C. psl.* complex were aligned with those of the 34 strains and outgroups using MAFFT ver. 6 (Katoh and Toh 2008). Gaps were omitted from the aligned sequences for phylogenetic analysis. For the Bayesian analysis, we applied an HKY + G model, selected by a hierarchical likelihood ratio test (hLRT) using MrModeltest v. 2.1 (Nylander 2004). Bayesian analysis was performed using MrBayes v. 3.1.2 (Ronquist and Huelsenbeck 2003), as described by Tsuchikane et al. (2012). Unweighted maximum-parsimony (MP), and neighbor-joining (NJ) analyses were performed as described by Tsuchikane et al. (2018b). The phylogenetic trees were rooted with sequences from *Closterium ehrenbergii* (DDBJ accession number: AY148821).

### Flow cytometry

Seeds of *Solanum lycopersicum* cv. Stupicke [standard for flow cytometry, 2C DNA content = 1.96 pg, (Doležel et al. 1992)] were kindly supplied by Dr. Jaroslav Doležel (Institute of Experimental Botany, Centre of Plant Structural and Functional Genomics, Olomouc, Czech Republic). DNA (1 pg) was converted into 978 Mbp (Doležel et al. 2003). The plants were grown from seeds and maintained at room temperature. Young leaves (0.1–0.05 g) were used as the standards. The quantity of nuclear DNA of the algae was estimated by flow cytometry using a CyFlow Ploidy Analyser “DAPI + PI” (Sysmex Corporation, Hyogo, Japan). CyStain PI Absolute P [for propidium iodide (PI) staining] (Sysmex Corporation) and CyStain UV Precise P [for 4′,6-diamidine-2-phenylindole (DAPI) staining] (Sysmex Corporation) were used as reagents for flow cytometric measurements according to the manufacturer’s instructions. Although PI and DAPI staining were performed using the same procedure, the composition of each reagent was different.

Algal vegetative cells in the logarithmic phase (5–10 days after inoculation) were collected at the beginning of the light period and counted. Thirty thousand cells were collected by centrifugation (1,450×*g*, 5 min), and the supernatant was discarded. To release the protoplasts, 100 μL of the enzymatic mixture of 0.5% Cellulase Onozuka R-10 (Yakult Pharmaceutical Industries Co. Ltd., Tokyo, Japan), 0.5% Macerozyme R-10 (Yakult), and 0.6 M Sorbitol (FUJIFILM Wako Pure Chemical Corporation, Osaka, Japan) dissolved in 30 mM 2-(*N*-morpholino) ethanesulfonic acid buffer (Wako) (pH 5.5) were added. The suspensions were rotated (20 rpm) for 1–3 h in the dark at 29°C. After incubation, the suspensions were centrifuged (18,000×*g*, 3 min), and the supernatant was discarded and replaced with 150 μL of Nuclei Extraction Buffer (Sysmex Corporation). The standard plant was chopped with a razor blade in Nuclei Extraction Buffer. The algal solution was added to the chopped standard and mixed, and the suspension of nuclei was filtered through a nylon mesh (20 μm nonsterile Cup Filcons, As One Corporation, Osaka, Japan) into a tube containing 1,600 μL of staining solution. Genome size was estimated based on the linear ratio between the 2C peak of the standard and the peak for each sample. In a single experiment, all strains were measured at least twice, usually thrice. Reproducibility was confirmed by performing the same experiment on different days. Data are shown for the second experiment. Two peaks were observed for the algae. The first peak was labeled 1C and the second 2C.

### Observation of chromosomes

Vegetative cells in the logarithmic phase (3–5 days after inoculation) were collected by centrifugation (1,450×*g*, 5 min), and the supernatant was discarded. The pellet was fixed in an ethanol mixture (ethanol/acetic acid, 3:1 v/v). Before observation, the cells were centrifuged at 300×*g* for 5 min and then repeatedly washed in Buffer B [10 mM Tris-HCl buffer (Wako), pH 7.6, containing 0.25 M sucrose (Wako), 1 mM di-sodium dihydrogen ethylenediaminetetraacetate dihydrate (EDTA, Nacalai Tesque Inc. Kyoto, Japan), 0.2 mM spermine (Wako), 0.4 mM spermidine (Wako), 0.25% NaCl (Wako), and 0.05% 2-mercaptoethanol (Wako)] (Ogawa et al. 1980). The nuclei were stained with 1 μg/mL DAPI (Nacalai), 10 ng/mL DABCO (Wako), and 0.05% Triton X-100 (Sigma-Aldrich, St. Louis, MO, USA) in Buffer B for 1 min, and observed concurrently. Fluorescent images were observed using an Olympus FV1200 confocal laser scanning microscope (filter set: U-FUW, Excitation Filter: 340–390, Dichroic Mirror: 410, emission filter: 420IF, Olympus, Tokyo, Japan). Each sample was recorded as a confocal Z-stack comprising approximately 70–78 optical slices of 0.04–0.05 μm thickness. Images were recorded using an FV10-ASW (Olympus). Each file consists of a confocal Z-stack comprising 70–78 individual images.

### Cell size measurement and statistical analysis

Algal vegetative cells in the logarithmic phase (5–10 days after inoculation) were collected at the beginning of the light period and counted. Thirty thousand cells were collected by centrifugation (1,450×*g*, 5 min), and the supernatant was discarded. The morphological features (length, width, and area of cells) were measured using ImageJ (Schneider et al. 2012). Regression analyses and t-tests were performed using Microsoft Excel for Mac 16.64.

## RESULTS

### Phylogenetic relationship between homothallic and heterothallic strains

In total 13 clones of the *C. psl.* complex were obtained from a wide geographical area (Table 1). The Hana, Naga37s-1, Shima19-9, Biwa5-3, Izu12-7-4, Mie54-1, Miya184-1, Naga56-6, Oki21-8, Yama58-4, and Ho36-3 strains were isolated and established from paddy fields. Zygospore formation was observed within clones derived from one cell (Intracrossing, Table S1). We therefore treated these strains as homothallic. The Oda12-4-1 and Oki23-2 strains were isolated and established from an agricultural water channel (waterways that supply water to paddy fields) in Kanagawa and Okinawa, respectively. Zygospore formation was not observed during intracrossing in these strains (Tables S2 and S3). Additionally, zygospore formation was observed in the mixture of Oda12-4-1 and mt^+^ cells of mating group II-A (Table S2) and in that of Oki23-2 and mt^−^ cells of mating group II-B (Table S3). We therefore treated these as heterothallic.

A Bayesian phylogenetic tree was constructed with an alignment of 376 bp combining the 1506 group I intron sequences from the *C. psl*. complex (Fig. 1). The sequences of the group I introns of Oda12-4-1 and Oki23-2 were identical to that of NIES-62 (mating group II-A) and of NIES-60 (mating group II-B), respectively. The monophyly of homothallic strains (homothallic clade A) of Naga37s-1, Shima19-9, and Hana was supported by high bootstrap values (100 and 89% in NJ and MP analyses, respectively) and a Bayesian posterior probability of 0.99 (Fig. 1). The heterothallic mating group II-A and these homothallic strains formed a monophyletic unit (Fig. 1, Clade A), despite the low Bayesian posterior probability and low bootstrap values for the NJ and MP analyses. The monophyly of homothallic strains (homothallic clade B) of NIES-2666, Biwa5-3, Izu12-7-4, Mie54-1, Miya184-1, Naga56-6, Oki21-8, and Yama58-4 was supported by high bootstrap values (100 and 94% in NJ and MP analyses, respectively) and a Bayesian posterior probability of 0.98 (Fig. 1). The heterothallic mating group II-B and these homothallic strains formed a monophyletic unit (Fig. 1, Clade B), albeit with relatively low statistical support (0.93 Bayesian posterior probability; bootstrap values of 60 and 64% in NJ and MP analyses, respectively). The monophyly of mating group II-C and the homothallic strain Ho36-3 (homothallic clade C) was supported by high bootstrap values (100 and 100% for NJ and MP analyses, respectively) and a Bayesian posterior probability of 0.99 (Fig. 1, Clade C). Our homothallic strains consisted of three clades (homothallic strains of clades A, B, and C). Clades containing homothallic and heterothallic strains were identified.

**Fig. 1.**
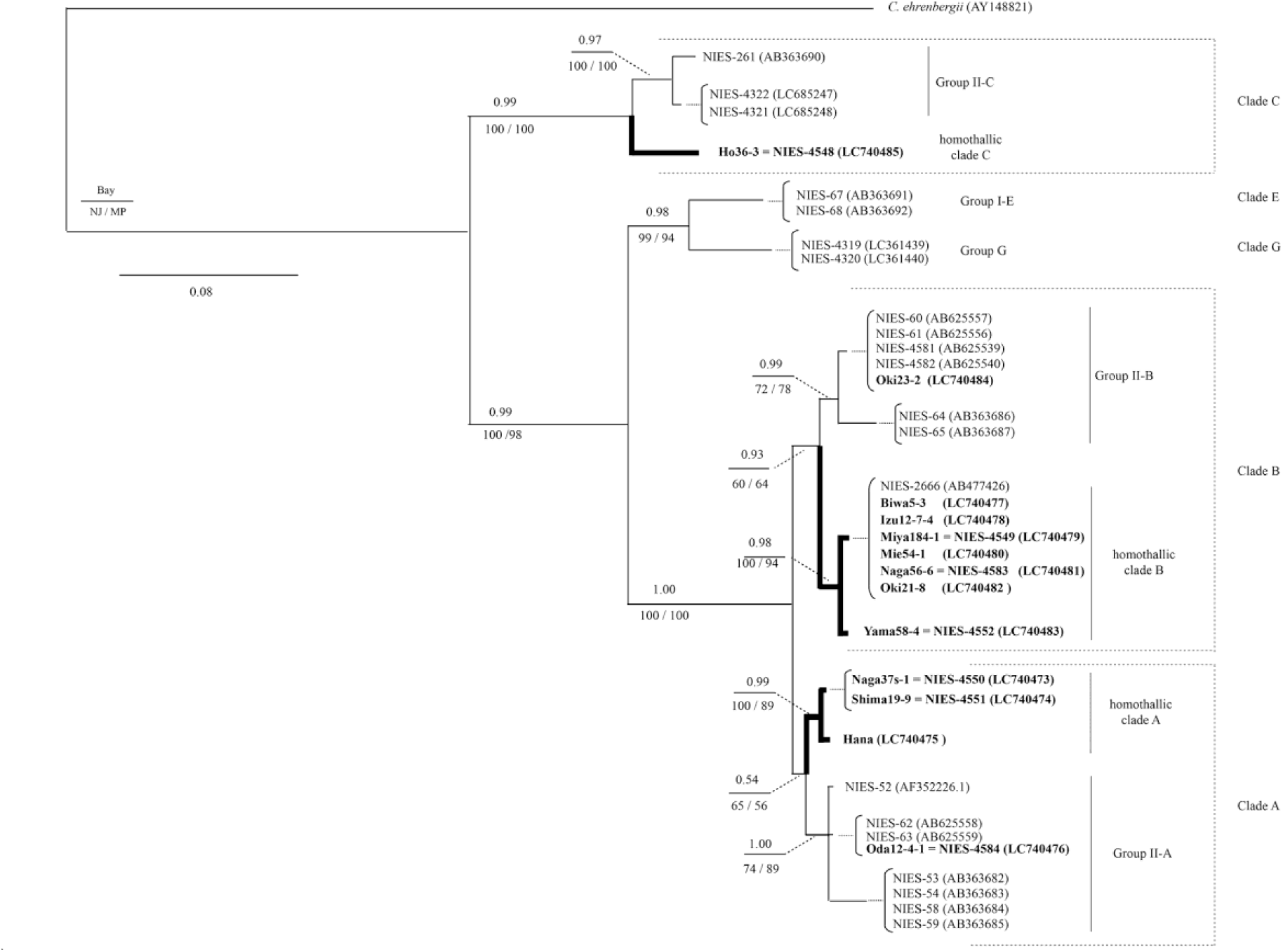
Bayesian phylogenetic tree based on 376 bp of the combined aligned 1506 group I intron. *Closterium* strains with identical sequences were treated as a single operational taxonomic unit. Numbers indicate posterior probabilities from the Bayesian analysis (Bay.; top), bootstrap values from neighbor-joining (NJ; bottom left) and maximum-parsimony (MP; bottom right) analyses. Branch lengths represent nucleotide substitutions per site. Bold branches indicate homothallic strains. The strains for which 1506 group I intron sequences were determined in this study are indicated in bold.

In summary, we identified three clades of homothallic strains, and each of them formed a clade with heterothallic strains, indicating at least two transitions between heterothallism and homothallism in the *C. psl* complex: one in clade C, and at least one in clades A and B.

### Measurement of genome size using flow cytometry

In the flow cytometric measurements after PI staining, the cells incubated under a 16 h light/8 h dark photoperiod were collected at the beginning of the light period (Fig. 2a, 0 h). The histogram usually showed two peaks in many strains (Fig. S1). In contrast, the histogram showed one peak only in NIES-53 (Fig. S2a and b). Flow cytometric measurements were tested using cells collected at various sampling points (Fig. 2a). NIES-68 and NIES-4321 showed two distinct peaks from the cells collected during the light or dark period, respectively (Fig. 2b and c). Although NIES-53 collected during the light period showed only one peak, two peaks were observed for the cells collected during the dark period (Figs 2d and S2). The values of the second peak were approximately twice those of the first peak (Fig. S3a). Therefore, the first peak was interpreted to represent vegetative haploid cells in the G1 phase (1C) and the second peak in the G2 phase (2C).

**Fig. 2.**
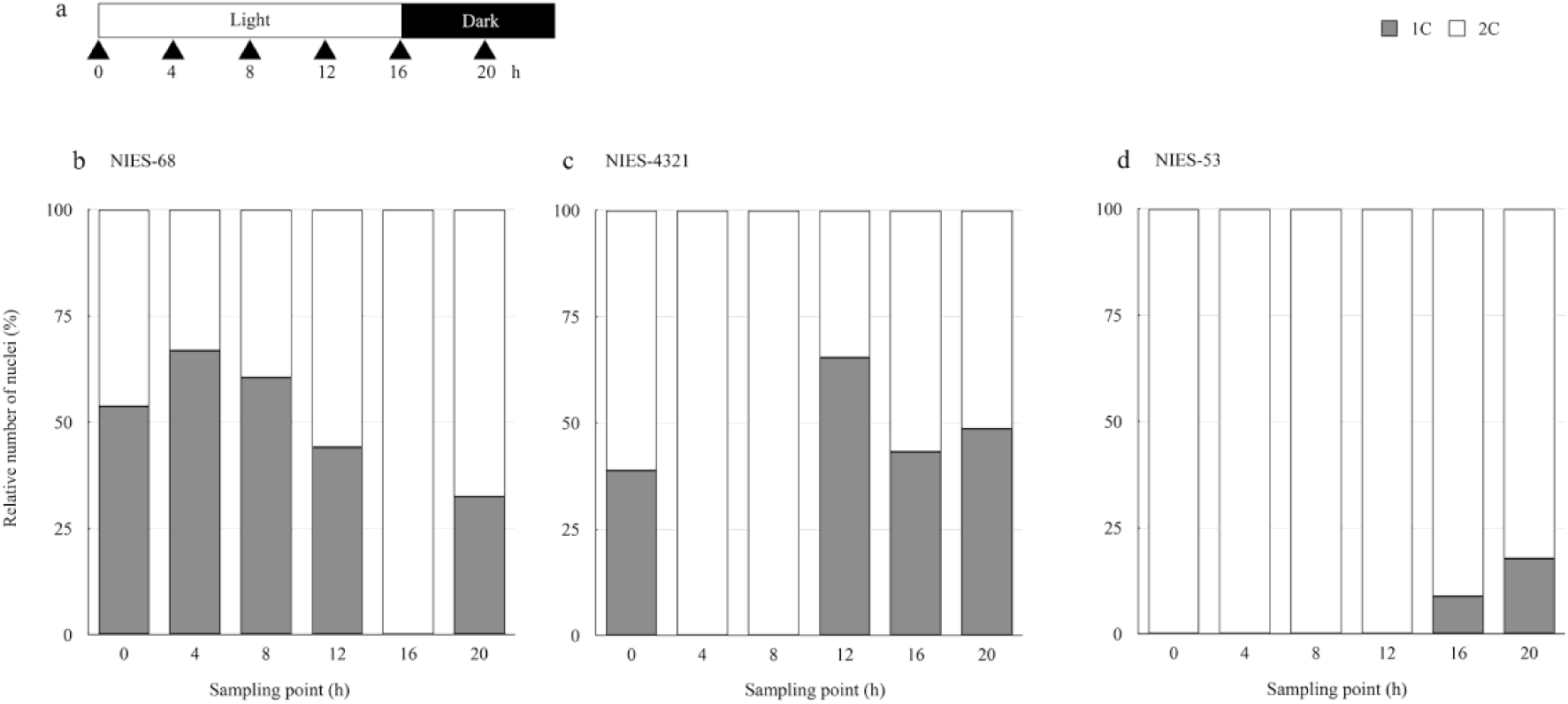
Ratio of 1C to 2C peaks at each sampling point under light and dark cycles. The number of nuclei at the peaks located at 1C (first peak, filled boxes) and 2C (second peak, open boxes) was measured. (a) The cells obtained at each sampling point under light and dark cycles were subjected to flow cytometry analysis using PI staining. (b) The sample volume for NIES-68 was 100 μL each. (c) The sample volume for NIES-4321 was 100 μL each. (d) The sample volume for NIES-53 was 250 μL each. Two experiments were performed at separate times, and similar results were obtained.

In total, the genome size was determined for six homothallic and 13 heterothallic strains (Fig. 3). High variability in genome size was detected between different strains of the *C. psl.* complex. The size of the 1C genome varied from 0.53 to 1.42 Gbp. Genome assemblies of the *C. psl.* complex were generated using a long-read (Pacbio) sequencing strategy, culminating from 0.33 to 0.98 Gbp of total contig length for the *C. psl.* complex (Sekimoto et al. 2023, Kawaguchi et al. 2023). The genome size was also estimated by k-mer analysis in these studies, ranging from 0.36 to 1.19 Gbp (Fig. 4a) (Sekimoto et al. 2023, Kawaguchi et al. 2023). We found that flow cytometry-based estimates were significantly correlated with both k-mer-based and assembly-based estimates (Fig. 4; *P* < 0.05). We note that assembly-based estimates were generally smaller than flow cytometry-based values, as reported in a wide range of species (Elliott and Gregory 2015). K-mer analysis also provided smaller genome size estimates, which can be variable depending on the coverage threshold (Sekimoto et al. 2023, Kawaguchi et al. 2023).

**Fig. 3.**
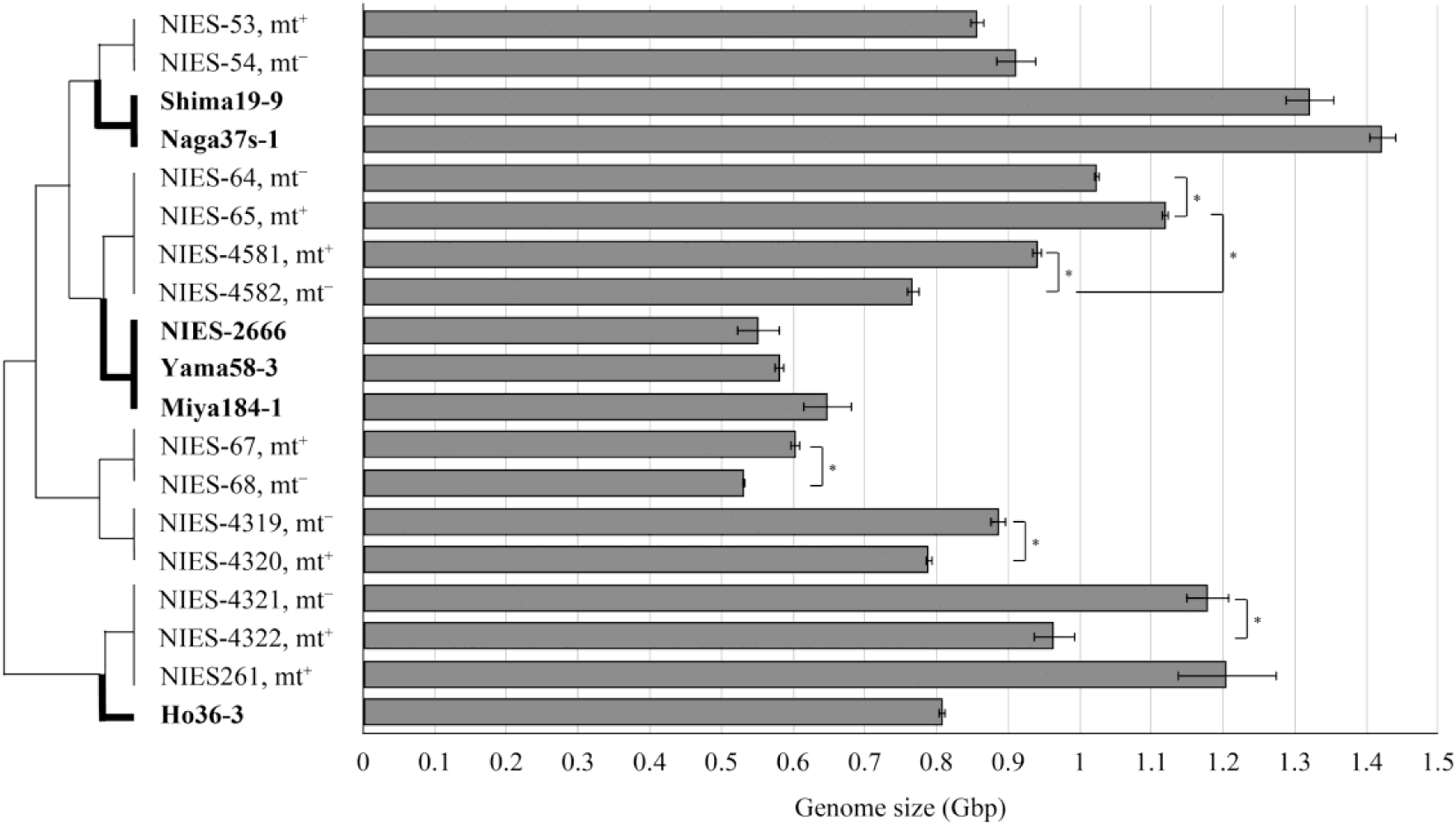
Genome size of the *C. psl.* complex. 1C genome size was measured using PI-stained samples (right). The phylogenetic relationship of each strain is shown (left). Bold branches indicate homothallic strains. *S. lycopersicum* cv. Stupicke serves as a reference standard. Horizontal lines depict standard errors (*n* = 3). *, t-test indicates significant difference (*P* < 0.05).

**Fig. 4.**
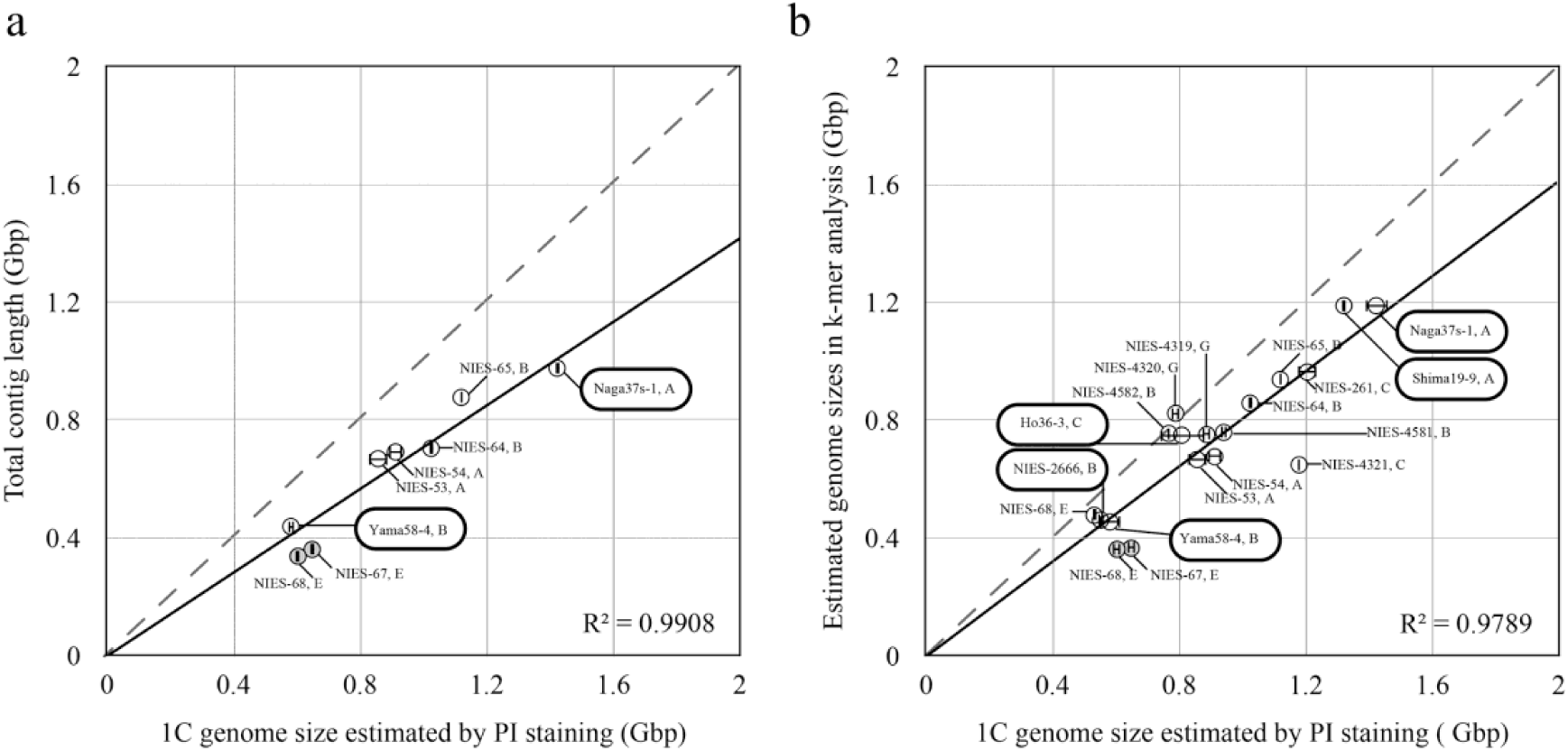
Comparison of genome size estimates between flow cytometry-based methods and assembly-based (a) or k-mer-based methods (b). Data reported by Kawaguchi et al. 2023 (open circles) and Sekimoto et al. 2023 (filled circles) were used. The dotted line indicates the auxiliary line (*y* = *x*). The bold round box indicates homothallic strains. Horizontal lines depict standard errors (*n* = 3). Significant correlations are indicated by solid lines (*P* < 0.05).

Significant differences in genome size were observed between mt^+^ and mt^−^ cells capable of zygospore formation in mating groups II-B, II-C, I-E, and II-G (Fig. 3). These differences varied between the strains from 8.5 to 31.5% (Table S4). We also found that mating systems (homothallism or heterothallism) and genome size were not significantly associated (P = 0.898). The genome sizes of homothallic strains Shima19-9 and Naga37s-1 in clade A were larger than those of heterothallic strains NIES-53 and NIES-54 (mating group II-A) (Fig. 3). By contrast, in clade B, the genome sizes of homothallic strains NIES-2666, Yama58-3, and Miya184-1 were smaller than those of heterothallic strains NIES-64, NIES-65, NIES-4581, and NIES-4582 (mating groups II-B). Therefore, we did not observe consistent trends of genome size changes in independently evolved homothallic strains. In clade A, homothallic strains had larger genome size but the sizes were less than twice those of heterothallic strains, suggesting that genome size differences between homothallic and heterothallic strains are not explained by simple polyploidization.

The genome sizes of 10 strains were also estimated using flow cytometry after DAPI staining. The genome sizes estimated with DAPI staining were approximately half those estimated with PI staining (Fig. 5, Fig. S3b). These differences varied between the strains from 41.8 to 65.2% (Table S5). Because DAPI binds preferentially to AT-rich regions, as discussed later, the PI-based estimates of genome size were used for further analyses.

**Fig. 5.**
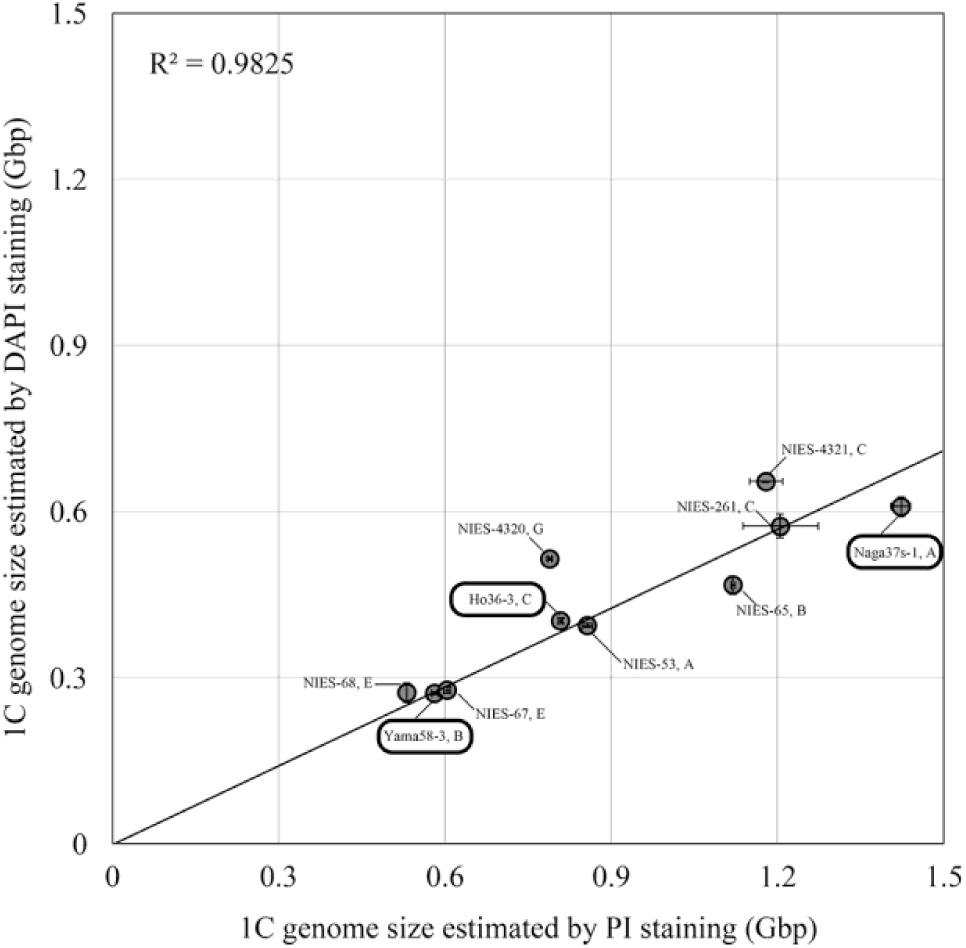
Relationship between 1C genome size measured in PI-stained samples and that in DAPI-stained samples. Horizontal and vertical lines depict standard errors (*n* = 3). Significant correlations are indicated by solid lines (*P* < 0.05). The bold round box indicates homothallic strains.

### Comparison of genome size and morphological features

The morphological features (cell size: length, width, and area of cells) of six homothallic and 13 heterothallic strains (Fig. S4) were measured and analyzed (Fig. S5, area). Morphological features between homothallic and heterothallic strains were compared, and no morphological features specific for mating systems were present (t-test, *P* > 0.05; Fig. 6a). Genome size was positively correlated with the cell size, and cell width showed a particularly strong correlation (Fig. 6b–d; area: *P* = 0.04099, length: *P* = 0.07388, width: *P* = 0.00032).

**Fig. 6.**
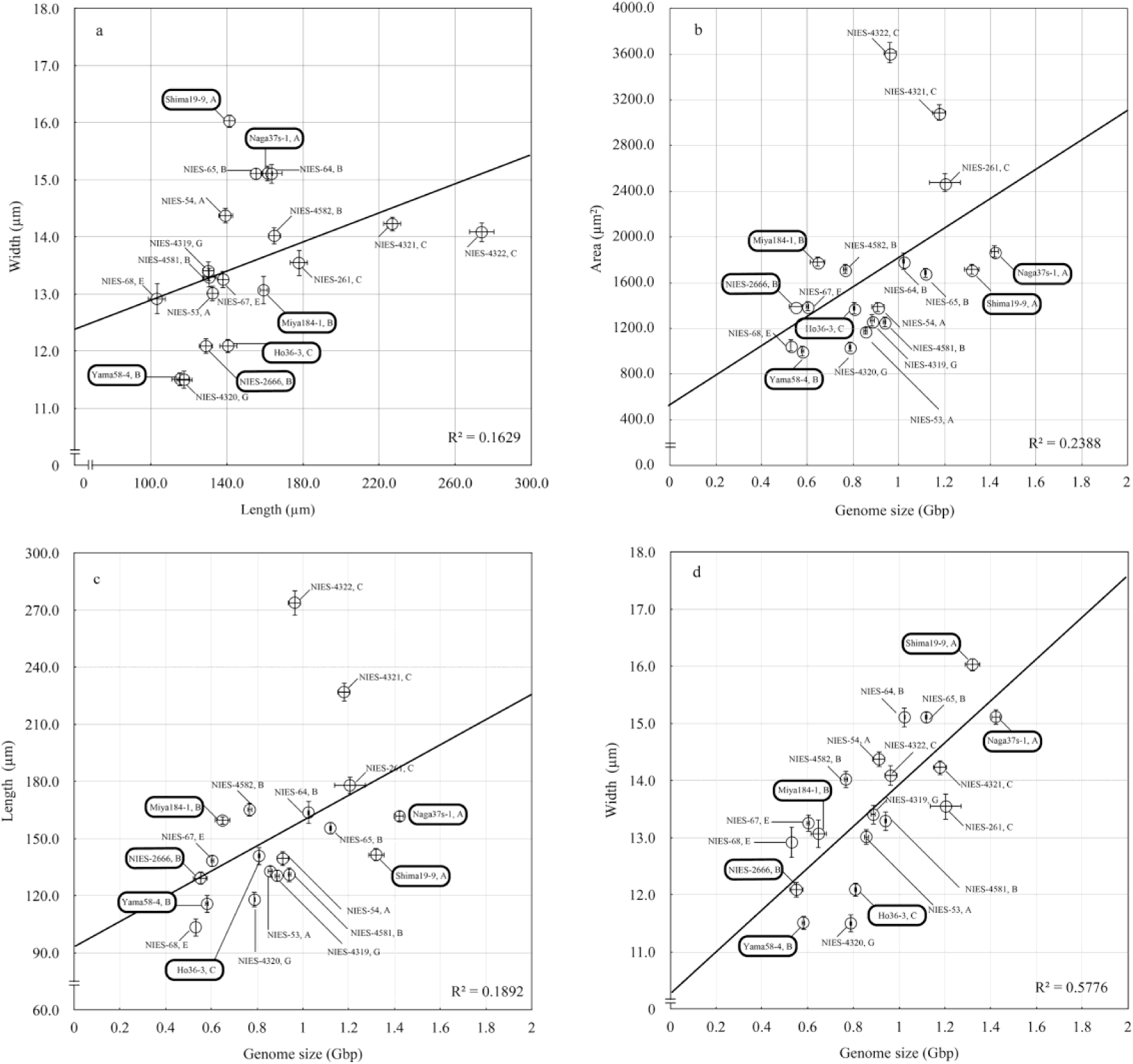
Correlation of genome size and morphometric parameters. (a) Variation in width and length between the strains. Mean values of each strain in width and length are shown. (b) Relationship between area and the genome size. (c) Relationship between length and genome size. (d) Relationship between width and genome size. Horizontal and vertical lines depict standard errors (*n* = 30). 1C genome size measured using PI-stained samples is shown. Strain and clade names are shown. The bold round box indicates homothallic strains. A regression line by linear model is indicated with a solid line (area: *P* = 0.04099, length: *P* = 0.07388, width: *P* = 0.00032).

### Comparison of genome size and chromosome number

The numbers of chromosomes in homothallic (Naga37s-1 and Yama58-4) and heterothallic (NIES-53, NIES-54, NIES-64, and NIES-65) strains were compared. The number of chromosomes was not counted exactly with high confidence because of the difficulty in spreading the chromosomes and distinguishing between small dot-like chromosomes without clear centromere structures (Godward, 1966) and because of background noise (Fig. S6). Nevertheless, the number of chromosomes in the two homothallic strains and four heterothallic strains were determined through analyses of confocal image stacks (Fig. 7a). Clearly different numbers of chromosomes were reproducibly obtained for different strains. The number of chromosomes varied from 26.6±1.31 to 87.7±5.55 (Fig. 7a). A significant correlation was found between the genome size and estimated number of chromosomes in the two homothallic (homothallic clades A and B) and four heterothallic strains (mating groups II-A and II-B) (Fig. 6a; R^2^ = 0.6855, *P* < 0.05). The mean chromosome sizes calculated as (genome size) / (chromosome number) are presented in Fig. 7b. In two clades, namely II-A and II-B with respective associated homothallic strains, the mean chromosome size in the homothallic strain was lower than for the related heterothallic strains.

**Fig. 7.**
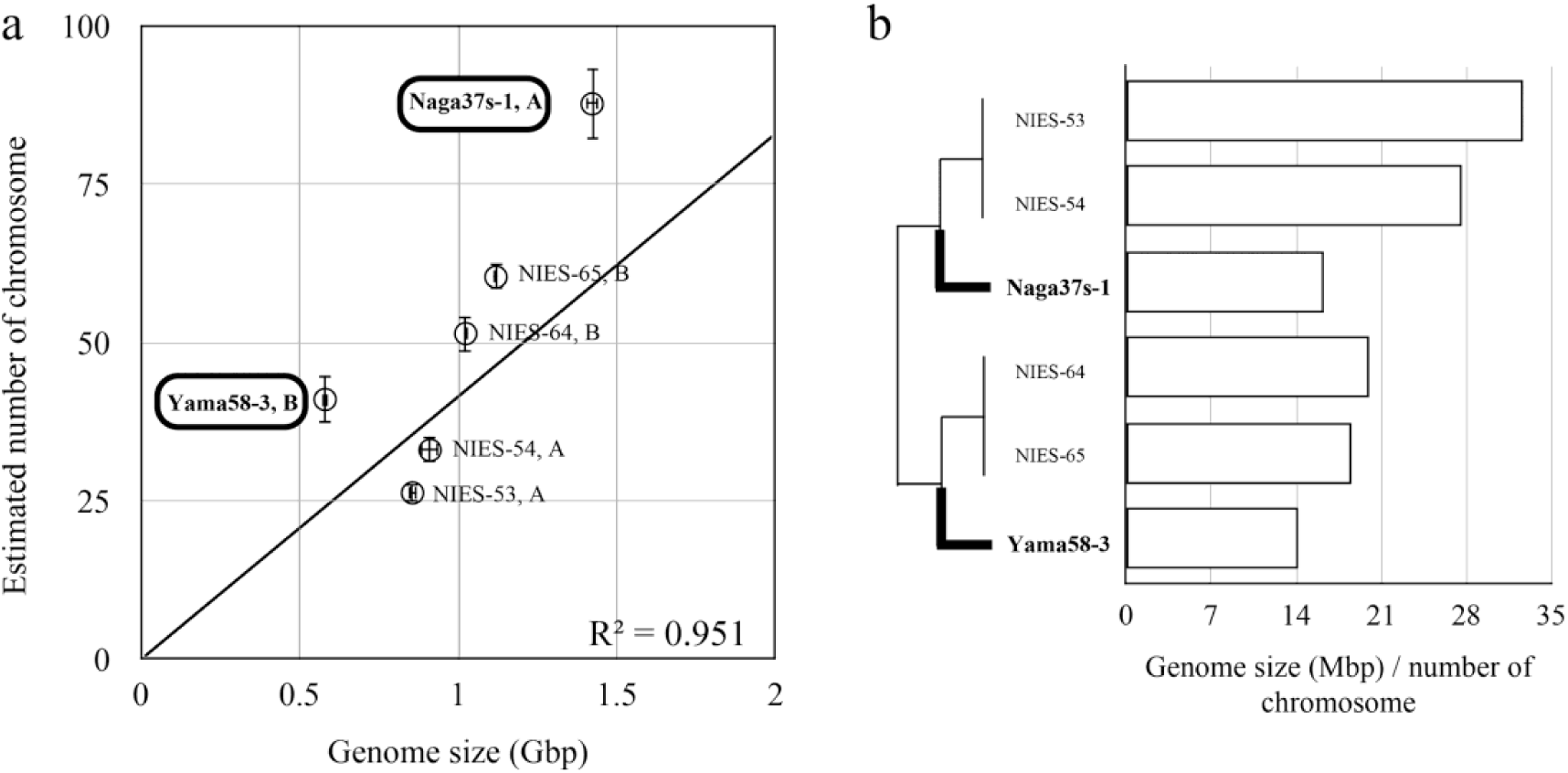
Correlation of 1C genome size and chromosome data. (a) The significant correlation of genome size and estimated number of chromosomes is indicated by solid lines (*P* < 0.05). The bold round box indicates homothallic strains. Horizontal lines depict standard errors of 1C genome size (*n* = 3). Vertical lines depict standard errors of chromosome number (*n* = 3). (b) Ratio of genome size to number of chromosomes (right). The phylogenetic relationship of each strain is shown (left). Bold branches indicate homothallic strains.

## DISCUSSION

### Evolution of homothallic and heterothallic strains

In this study, we investigated the phylogenetic relationship of the heterothallic and homothallic strains in the *C. psl.* complex (Fig. 1). Both homothallic and heterothallic strains were found in all of the clades A, B, and C, and clades A and B formed a monophyletic unit, albeit with relatively low Bayesian posterior probability and low bootstrap values in the NJ and MP analyses. These results suggest that transitions between homothallism and heterothallism occurred at least twice in the *C. psl.* complex. Multiple transitions between homothallism and heterothallism are also supported by a single nucleotide polymorphism-based phylogeny in a recent report (Kawaguchi et al. 2023). This system will be useful to further investigate common biological features underlying mating system transitions by comparing independently evolved homothallic and heterothallic strains.

Homothallic strains can rapidly form dormant spores (zygospores) between sister gametangial cells that have just divided. Such a trait may be adaptive under an environment with fluctuating water availability because they can form dormant spores even in a clonal population. Consistent with this hypothesis, homothallic strains were isolated from Japanese paddy fields, which are drained annually (Hendrayanti et al. 2004, Tsuchikane et al. 2010), while heterothallic strains were mainly isolated from waterways, lakes, and ponds (Table 1; Kobayashi et al. 2022, Tsuchikane et al. 2018b). This idea could be tested through a field survey on a larger scale, by employing a high-throughput identification method such as metagenomic analysis.

### Genome size measurement using flow cytometry

Flow cytometry has been used extensively to estimate genome size in land plants, but its application has been limited in zygnematophycean algae. Here, we discuss technical aspects of using flow cytometry for zygnematophycean algae.

Cells in the G2 phase, where DNA is replicated, were always detected in the *C. psl.* complex (Fig. 2). In contrast, cells in the G1 phase were minor, depending on the strain (Fig. 2). Therefore, genome size measurement using flow cytometry in a *Closterium* strain for the first time should detect two peaks and allow the correct value of the 1C peak to be assigned. In *Physcomitrella* haploid gametophore leaves, 1C peaks of cells in the G1 phase and 2C peaks of cells in the late S phase are observed (Ishikawa et al. 2011). Therefore, there may be haploid organisms other than *Closterium* that require observation of the 1C and 2C peaks.

The genome size determined for DAPI-stained samples was approximately half of the genome size determined for PI-stained samples (Fig. 5). The AT content of the *S. lycopersicum* genome is 66% (Sato et al. 2012). In contrast, the AT contents of NIES-67 and NIES-68 were 43.9 and 44.2%, respectively (Sekimoto et al. 2023). When using DAPI-stained samples, the genome size cannot be measured accurately if organisms with different AT% are used as controls. Additionally, DAPI has affinity for the AATT site in the base sequence, and the affinity decreases in the order: AATT >> TAAT ≈ ATAT > TATA ≍ TTAA (Breusegem et al. 2002). The difference in the distribution of AATT sites between strains may result in uneven gaps between the genome sizes of DAPI-stained and PI-stained samples between strains (Table S5). Indeed, PI-stained samples are used in land plants and zygnematophycean algae for genome size estimation (Doležel et al. 1998, Poulíèková et al. 2014, Mazalová et al. 2011). Future genome size measurements of zygnematophycean or other algae should confirm the two peaks and use PI-stained samples for flow cytometry.

### Extensive genome size variation and its correlation with other factors

In this study, we identified more than twofold genome size variation in the *C. psl* complex, ranging from 0.53 to 1.42 Gbp, as revealed by flow cytometry analysis. In this section, we discuss what factors are correlated with this genome size variation, specifically, mating types of heterothallic strains, chromosome numbers, cell size, and mating systems.

We found that, in heterothallic strains, genome size was not significantly correlated with mating types (mt^+^ and mt^−^) based on the presence or absence of the mating-type determining gene, *CpMinus1* (Tsuchikane and Sekimoto 2019, Sekimoto et al. 2023). The mating groups G and I-E were reproductively isolated from each other, although the sexual reaction and release of lone protoplasts from mt^−^ cells of mating group G were induced in the presence of mt^+^ cells of mating group I-E (Tsuchikane et al. 2018b). Mt^+^ cells showed a larger genome size than mt^−^ cells containing *CpMinus1* in mating group I-E, but mt^−^ cells containing an ortholog of *CpMinus1* showed a larger genome size than mt^+^ cells in mating group G (Figs 3 and S1), suggesting that genome size variation is not simply due to structural variation linked to the mating-type determining locus.

The observation of chromosome number revealed that genome-size variation is partly explained by the chromosome number variation between strains, given the positive correlation between them (Fig. 7a). Such chromosome number variation was also reported in *Closterium ehrenbergii* Meneghini. In sexually isolated mating groups of *C. ehrenbergii*, chromosome number was as diverse as 100–106 for mating group A, 97–104 for mating group B, and 105–127 for mating group C (Kasai and Ichimura 1984). A positive correlation between genome size and chromosome number was also observed in other zygnematophycean alga, *Micrasterias* (Poulíèková et al. 2014).

It is notable that extensive genome size variation was observed even between pairs that can form zygospores. NIES-64 and NIES-65 strains showed an 8.5% difference in genome size (Fig. 3 and Table S4), but zygospores were formed and F1 progenies were produced, both of which had been isolated from the same locality (Watanabe and Ichimura 1982, Kawaguchi et al. 2023). NIES-65 and NIES-4582 strains showed a 31.5% difference in genome size (Figs 3 and S1), but zygospores were formed between them (Tsuchikane et al. 2012). Similarly, hybrid zygospores were formed between homothallic strain NIES-2666 and heterothallic strain NIES-4581 (Tsuchikane et al. 2012), but the genome size difference was as much as 41.4% (Fig. 3 and Table S4). Future studies need to investigate whether F1 progenies are viable and whether postzygotic isolation is observed in this zygospore. Because we found that a large genome size variation within the mating group was partly due to the difference in chromosome number, there may be a mechanism tolerating chromosomal rearrangements during meiosis in the *C. psl.* complex.

Positive correlations were observed between the genome size and the morphological features representing cell size in *Closterium* (area: *P* = 0.04099, length: *P* = 0.07388, width: *P* = 0.00032). Correlation between genome size and cell size was also reported in *Micrasterias* (Poulíèková et al. 2014), which belongs to the same family Desmidiaceae as *Closterium* (Hess 2022). In *Closterium*, we found that correlation with genome size was strong in cell width but relatively weak in cell area and cell length. We speculate that a difference in cell cycles between strains explains this pattern. Each strain had a different cell cycle, based on the observation that the proportion of cells in G1 and G2 phases differed between strains (Fig. 2, 0 h). Given that cells harvested early in the light period were always used for cell size measurement, cells used in this study should be heterogeneous in terms of cell cycle. Cell width should be less influenced by cell division than cell length and area, possibly resulting in higher correlation with genome size. For precise comparison of the cell size, estimation of cell cycle state and sorting the data accordingly warrant consideration.

Although mating systems were not significantly correlated with genome size (t-test, *P* = 0.898), we found that, compared with heterothallic strains, homothallic strains generally showed a larger number of chromosomes relative to genome size, thus possibly having more fragmented chromosomes (Fig. 7b). In heterothallic populations, such chromosomal rearrangements are more likely to be eliminated by selection against pairing of different chromosomal structures during meiosis. However, in homothallic populations, zygospores can be formed from the conjugation of two clonal gametangial cells, allowing pairing of the same chromosomal structures. Therefore, homothallic populations may tolerate more chromosomal rearrangements including fragmentation suggested in this study. This scenario would be analogous to the argument that self-fertilizing populations are more prone to fix structural rearrangements in flowering plants (Wright et al. 2013).

### Diversity in chromosome number and genome size

Mechanisms of creating diversity in chromosome number and genome size in *Closterium* remain to be elucidated. A possible factor may be degradation of nuclei during meiosis. Two cells of different mating types germinate from a single zygospore after meiosis in *Closterium* (Kasai and Ichimura 1983, Hamada 1987). In anaphase II, the chromatids finally separate and become chromosomes in their own right; they are pulled to opposite poles in each of the two cells. In telophase II, the chromosomes gather in the nuclei, although the cells are not divided. Consequently, germinated cells have two nuclei in one cell and one of the two is disassembled in *Closterium* (Kasai and Ichimura 1983). If this degradation is insufficient, chromosomes that cannot be resolved will be retained and a part of the genome could be translocated to the nucleus, possibly resulting in increased genome size and chromosome numbers. In addition, Zygnematophycean algae are suggested to have holocentric chromosomes (Mandrioli and Manicardi 2020), which may stabilize chromosomal fragments and tolerate karyotype rearrangements because of their diffuse kinetochores. Other factors, such as transference of retrotransposons, which was observed in *Penium margaritaceum* (Jiao et al. 2020), and extensive gene duplication, may also be important to shape a large genome size variation in the *C. psl* complex (Kawaguchi et al. 2023). More detailed genome and chromosome analyses are required to understand how genome size diversity is generated and maintained.

## ACKNOWLEDGMENTS

This work was supported by the Grants-in-Aid for Scientific Research (nos. 25304012, 26650147, 18K06367, and 19K22446 to H.S., 19K22448 to T.N., H.S., and Y.T., 15H05237 to H.S. and Y.T., 16H04836, 16K02518, 18K19365, 20K21451, and 21H02549 to T.N. and H.S, and 22H05177 to T.N., 19K06827 to Y.T., 15K18583 and 17K15165 to TT) from the Japan Society for the Promotion of Science.

## ABBREVIATIONS

DAPI: 4′,6-diamidine-2-phenylindole;
MP: maximum parsimony
NJ: neighbor joining;
PI: propidium iodide

