## Supplementary Material for "DIVERSITY OF GENOME SIZE AND CHROMOSOME NUMBER IN HOMOTHALLIC AND HETEROTHALLIC STRAINS OF THE *CLOSTERIUM PERACEROSUM–STRIGOSUM–LITTORALE* COMPLEX (DESMIDIALES, ZYGNEMATOPHYCEAE, STREPTOPHYTA)"

Supplementary Tables

**Table S1.** Intracrosses of homothallic strains.

| Strain designation | Relative number of zygospores ± S.E. (%) | |
|---|---|---|
| Hana | Z | n = 2 |
| Naga37s-1 | 27.3 ± 1.2 | n = 3 |
| Shima19-9 | 76.4 ± 1.9 | n = 3 |
| Biwa5-3 | Z | n = 2 |
| Izu12-7-4 | Z | n = 2 |
| Mie54-1 | Z | n = 2 |
| Miya184-1 | Z | n = 2 |
| Naga56-6 | 36.0 | n = 2 |
| Oki21-8 | 14.7 | n = 2 |
| Yama58-4 | Z | n = 2 |
| Ho36-3 | 47.2 ± 1.9 | n = 3 |

Z, zygospores were reproducibly formed but the numbers were not counted.

**Table S2.** Intercrosses between strains of group II-A.

| | NIES-58 (het / –) | NIES-59 (het / +) | Oda12-4-1 |
|---|---|---|---|
| NIES-58 (het / –) | / | Z | / |
| NIES-59 (het / +) | | / | Z |
| Oda12-4-1 | | | / |

Z; zygospore formation was observed, /; no zygospore formation was observed.

**Table S3.** Intercrosses between strains of group II-B.

| | NIES-64 (het / –) | NIES-65 (het / +) | Oki23-2 |
|---|---|---|---|
| NIES-64 (het / –) | / | Z | Z |
| NIES-65 (het / +) | | / | / |
| Oki23-2 | | | / |

Z; zygospore formation was observed, /; no zygospore formation was observed.

**Table S4.** Differences in genome size between zygospore-forming pairs.

| Mating group | NIES No. | Mating type | 1C genome size ± SE (Gbp) | Mating type difference (%) |
|---|---|---|---|---|
| II-A | NIES-53 | $mt^+$ | 0.857 ± 0.010 | – |
| | NIES-54 | $mt^-$ | 0.912 ± 0.028 | 6.1 |
| II-B | NIES-65 | $mt^+$ | 1.119 ± 0.005 | 8.5 [a] |
| | NIES-64 | $mt^-$ | 1.024 ± 0.005 | – |
| II-B | NIES-4581 | $mt^+$ | 0.941 ± 0.006 | 18.5 [a] |
| | NIES-4582 | $mt^-$ | 0.767 ± 0.009 | – |
| II-B | NIES-65 | $mt^+$ | 1.119 ± 0.005 | 31.5[a] |
| | NIES-4582 | $mt^-$ | 0.767 ± 0.009 | – |
| II-B | NIES-4581 | $mt^+$ | 0.941 ± 0.006 | 41.4 [a] |
| hom | NIES-2666 | n.a. | 0.551 ± 0.003 | – |
| II-C | NIES-4322 | $mt^+$ | 0.964 ± 0.029 | – |
| | NIES-4321 | $mt^-$ | 1.179 ± 0.030 | 18.2 [a] |
| I-E | NIES-67 | $mt^+$ | 0.604 ± 0.007 [a] | 12.0 [a] |
| | NIES-68 | $mt^-$ | 0.531 ± 0.002 | – |
| G | NIES-4320 | $mt^+$ | 0.789 ± 0.005 | – |
| | NIES-4319 | $mt^-$ | 0.887 ± 0.011 | 11.1 [a] |

[a]t-test found significant difference ($P < 0.05$)

**Table S5.** Ratio of 1C genome size with DAPI staining to 1C genome size with PI staining.

| | % ± SE |
|---|---|
| NIES-53 | 46.0 ± 0.6 |
| Naga37s-1 | 42.4 ± 1.6 |
| NIES-65 | 41.8 ± 1.9 |
| Yama58-3 | 46.7 ± 1.3 |
| NIES-67 | 46.8 ± 1.5 |
| NIES-68 | 51.3 ± 3.4 |
| NIES-4320 | 65.2 ± 0.7 |
| NIES-4321 | 55.5 ± 1.6 |
| NIES261 | 51.7 ± 1.8 |
| Ho36-3 | 49.1 ± 0.1 |

Supplementary Figures

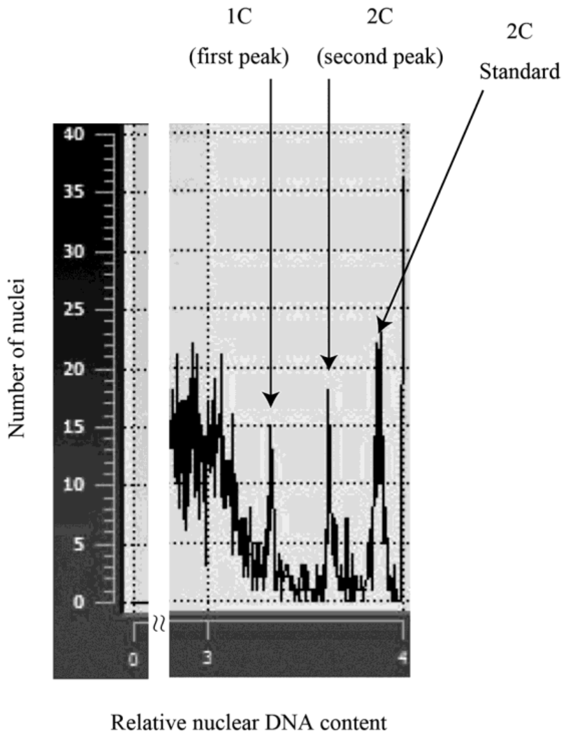


**Fig. S1.** Histogram of relative nuclear DNA content of the *C. psl.* complex (NIES-68) and standard. Example of the two peak (first and second peak) pattern is shown and treated as 1C and 2C, respectively. Standard (2C) is *Solanum lycopersicum*.

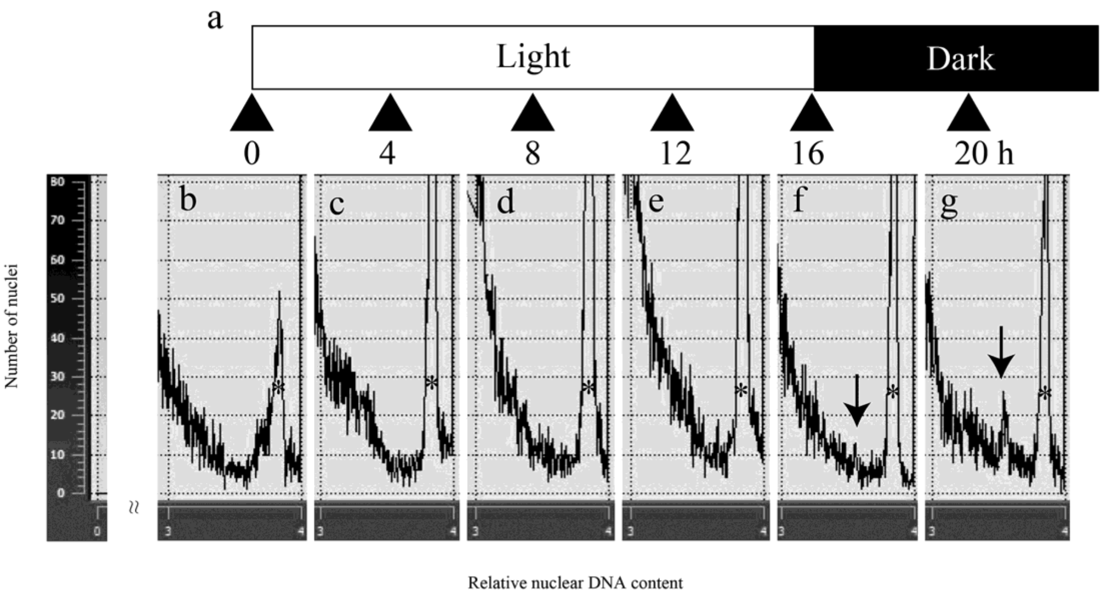


**Fig. S2.** Histogram of relative nuclear DNA content of NIES-53 obtained at each sampling point under light and dark cycles. The cells obtained at each sampling point under light and dark cycles were subjected to flow cytometry analysis using PI staining (a). Histogram of cells 0 (b), 4 (c), 8 (d), 12 (e), 16 (f), and 20 h (g) after the light period started. The sample volume was 250 μL each. Arrows indicate the 1C peak of NIES-53. Asterisks indicate the 2C peak of NIES-53.

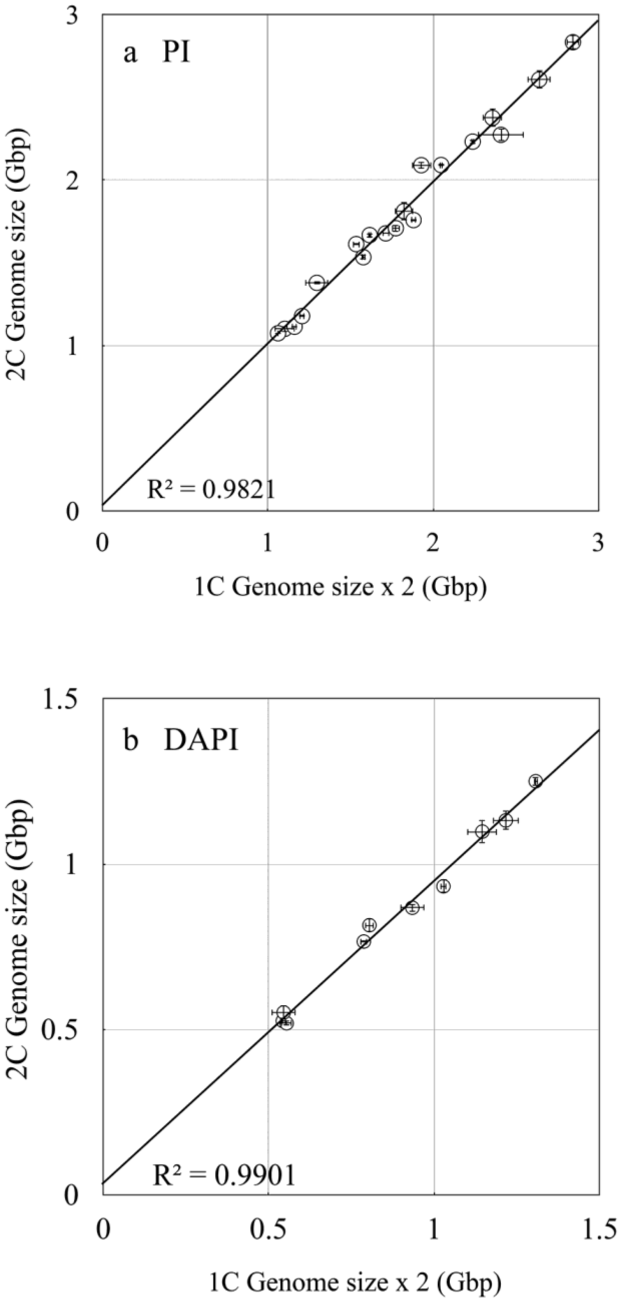


**Fig. S3.** Comparison of 2C genome size and doubled 1C genome size. Genome size obtained using PI-stained (a) and DAPI-stained (b) samples. Significant correlations are indicated by solid lines ($P < 0.05$).

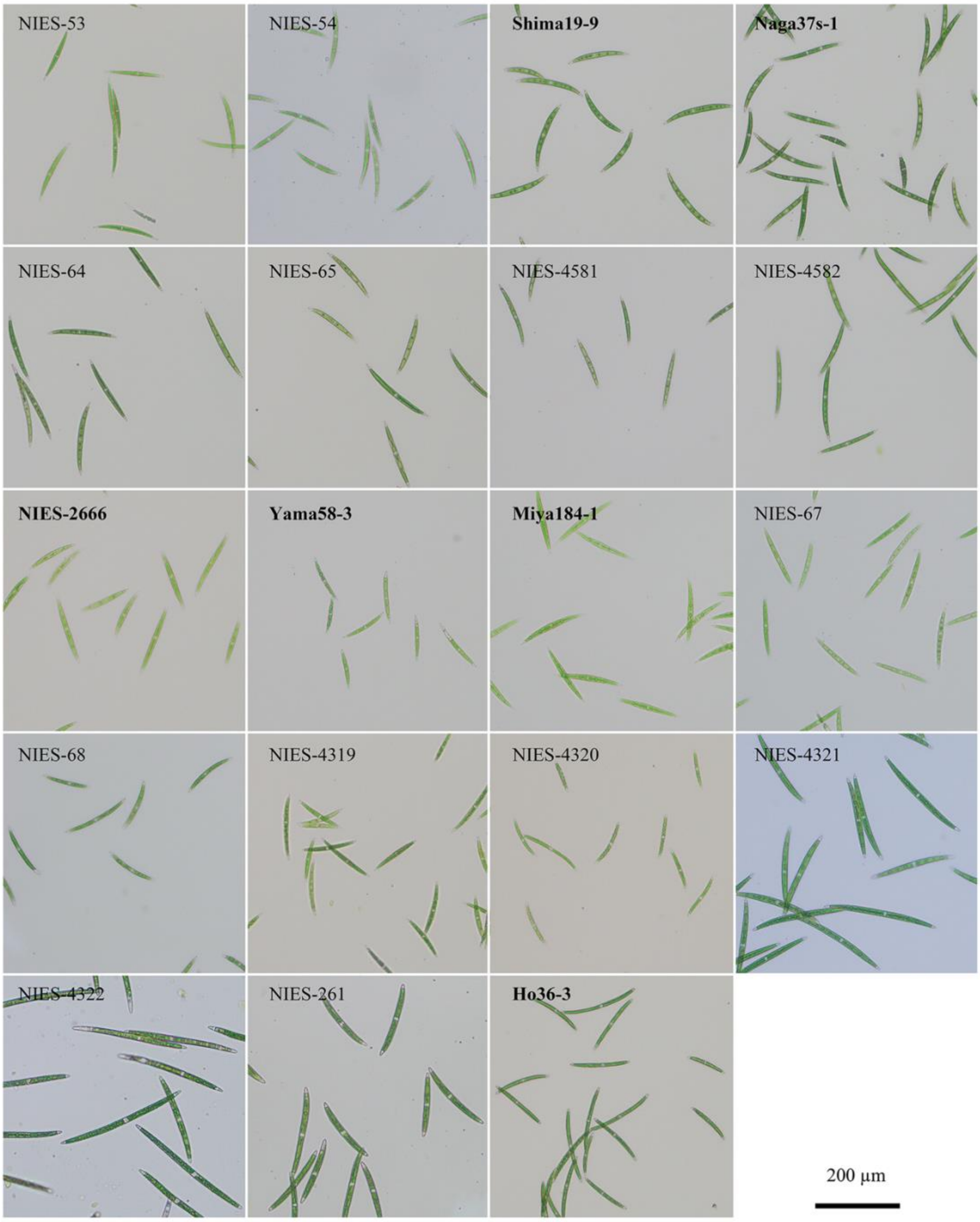


**Fig. S4.** Photographs of vegetative cells. Scale bar 200 μm. Homothallic strains are indicated in bold.

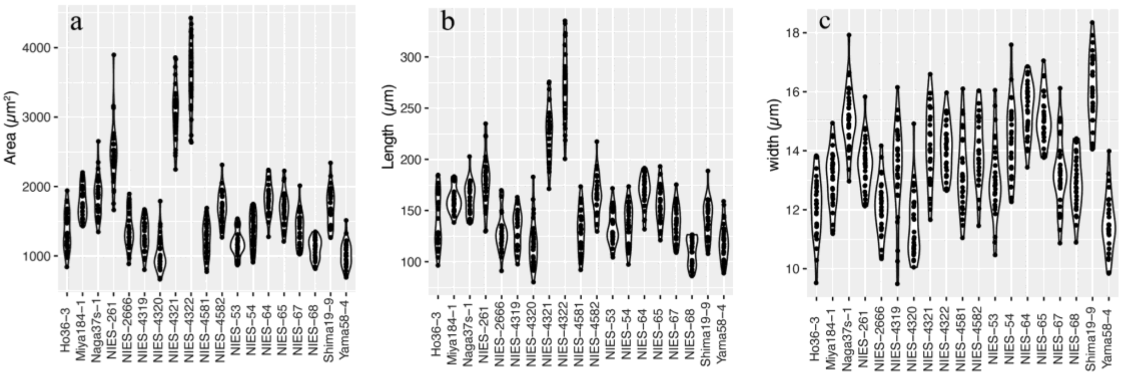


**Fig. S5.** Violin plots of cell size: (a) area, (b) length, (c) width.

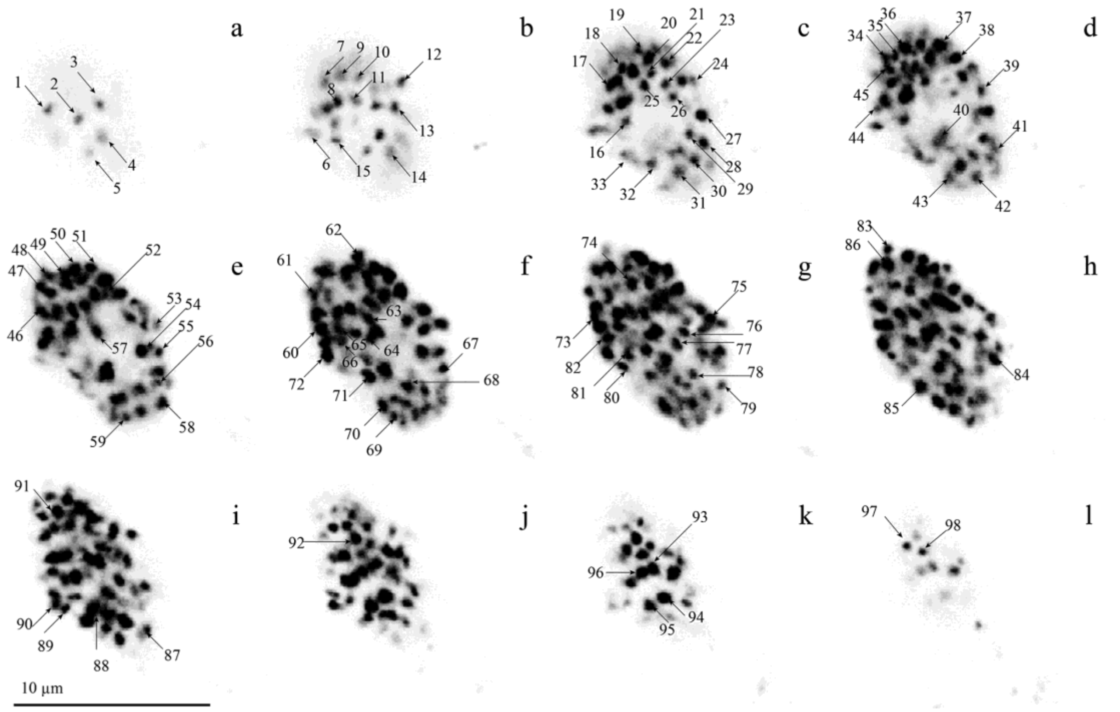


**Fig. S6.** Observation of chromosomes. Vegetative cells were DAPI stained. Chromosome number was measured in each plane on the *x*-axis. Photographs show the nuclei of Naga37s-1. Arrows indicate chromosomes. Scale bar 10 μm.
